# TENT-5 regulates the expression of male-specific genes in *Caenorhabditis elegans*

**DOI:** 10.1101/2024.06.18.599341

**Authors:** Zuzanna Mackiewicz, Vladyslava Liudkovska, Andrzej Dziembowski

## Abstract

Polyadenylation is an important post-transcriptional process that governs mRNA stability and expression. Advancements in direct RNA sequencing in recent years have clarified many aspects of this intricate regulation, revealing the influence of various factors. Here, we used Nanopore Direct RNA Sequencing to investigate the association between genome-wide mRNA poly(A) tail profiles and sexual dimorphism in *Caenorhabditis elegans*. Our results demonstrate sex-dependent differences in both gene expression and poly(A) tail metabolism. Notably, we discovered that cytoplasmic poly(A) polymerase TENT-5 regulates multiple male-specific transcripts, predominantly encoding putative seminal fluid components with predicted extracellular localization. TENT-5 expression in male-specific tissues, such as seminal vesicle and vas deference, corroborates its functional significance. Intriguingly, despite extensive TENT-5-mediated polyadenylation of male-specific transcripts, males devoid of TENT-5 show no abnormalities in mating behavior, sperm morphology, or fertility. Our findings suggest that TENT-5 plays a role in regulating sex-related processes in males, although the physiological consequences remain to be fully elucidated.

## Introduction

Polyadenylation is a crucial post-transcriptional process that modifies RNA species by adding adenosines to their 3′ ends, thereby influencing RNA stability and function within the cell. This modification primarily occurs in the nucleus, catalyzed by canonical poly(A) polymerase, and is essential for mRNA biogenesis and subsequent export to the cytoplasm (1, 2). Upon reaching the cytoplasm, mRNAs associate with poly(A) binding proteins, which are critical for maintaining mRNA stability and enabling efficient protein synthesis (2, 3). During the later lifespan of mRNA, poly(A) tails undergo further modifications. Deadenylating enzymes, such as Ccr4-NOT or PAN2/3, gradually shorten poly(A) tails, directing mRNAs to degradation (2, 4, 5). However, poly(A) tails can also be re-adenylated in the cytoplasm by noncanonical poly(A) polymerases (ncPAPs), which are members of the terminal nucleotidyltransferase family of proteins (TENTs) (6, 7). Cytoplasmic polyadenylation enhances stability, promotes the expression of selected transcripts, and prevents their degradation, contributing to the complex dynamics of poly(A) tails and regulation of gene expression (6, 7).

Recent advances in RNA sequencing technologies have enabled more comprehensive studies of poly(A) tail metabolism (8). Genome-wide poly(A) tail profiling has been carried out for various organisms, including yeast (9, 10), nematodes (11–13), mice (10, 14–16), and humans (10, 12, 17). These studies revealed organism-specific characteristics of poly(A) tails and their dependence on multiple factors such as developmental stage, age, tissue localization, and response to environmental cues. While poly(A) tail length is often positively associated with mRNA half-life and translation efficiency, this relationship is not always consistent. During early embryogenesis, longer poly(A) tails enhance mRNA stability and translation (2, 10, 18).

However, in somatic cells, poly(A) tail length often shows a poor or even negative correlation with mRNA expression levels and translational activity (10–12). For instance, mRNAs encoding highly abundant housekeeping machinery, such as ribosomal proteins, typically have short tails. Conversely, longer tails are predominantly found on lowly expressed mRNAs, which often encode regulatory proteins (11, 19). Moreover, poly(A) tail dynamics is affected by additional layers of regulatory control, including poly(A)-modifying enzymes that influence numerous physiological processes. For example, cytoplasmic polyadenylation by GLD2 (Germ Line Development 2)/TENT2 is indispensable for proper germline development and fertility in nematodes (20–22) and flies (23, 24). In mammals, a similar role is observed for the recently discovered TENT5 ncPAPs (15). Additionally, members of the TENT5 family have been implicated in regulating humoral and innate immune responses by stabilizing mRNAs encoding immunoglobulins and secreted defense proteins (13, 16). Despite these findings, the high complexity of cytoplasmic polyadenylation requires further examination to elucidate the interplay between poly(A)-modifying enzymes, the mechanisms driving their substrate specificity, and their effects on various groups of transcripts during multiple physiological processes and conditions (6).

Here, we demonstrate that sexual dimorphism is a critical factor that strongly influences polyadenylation profiles. For our studies, we used *Caenorhabditis elegans*, which features two distinct sexes: self-fertile hermaphrodites (XX) and males (X0), comprising approximately 0, 05% of the population. Utilizing Oxford Nanopore Direct RNA Sequencing (DRS), the most reliable methodology for studying poly(A) tails, we uncovered profound differences in gene expression and poly(A) length distributions between hermaphrodites and males. Notably, we showed that TENT-5 ncPAP is responsible for the regulation of male-specific transcripts that mainly encode secreted seminal fluid components. Our work provides new and interesting insights into the polyadenylation process, expanding our knowledge of both TENT-5 functions and *C. elegans* male physiology and genetics.

## Methods

### C. elegans culture and growth conditions

The following *C. elegans* strains were used: wild type N2 Bristol, BS553 (*fog-2(oz40) V*), and DR94 (*unc-45(m94) III*), obtained from the Caenorhabditis Genetics Center (CGC); *tent-5(tm3504) I*, obtained from the National Bioresource Project of Japan (NBRP); and ADZ21 (*tent-5(rtt6[tent-5::gfp::3xflag]) I*), previously generated in our laboratory (13). All strains were cultured at 20°C on nematode growth medium (NGM) plates seeded with *E. coli* HB101 as a food source. Male enrichment of N2 and *tent-5(tm3504) C. elegans* cultures was achieved through mating, with one hermaphrodite and five males of the same genotype placed on plates with a drop of *E. coli* HB101. Successful mating resulted in up to 50% male enrichment.

### Preparation of samples for Direct RNA sequencing (DRS)

Wild-type or *tent-5(tm3504)* young adult males were hand-picked from the male-enriched plates and transferred to an Eppendorf tube containing 10 µl of 50 mM NaCl. The worm pellet was then resuspended in 200 µl of TRI Reagent (Sigma-Aldrich), vortexed for 15 min at room temperature, and stored at -80°C before RNA isolation. This process was repeated three times to obtain sufficient biological material. TRIzol samples from three different pickings were combined just before RNA isolation, resulting in approximately 300 male worms per sample. The wild-type hermaphrodite population was age-synchronized by bleaching gravid adults and seeding isolated embryos on plates. Worms were grown until they reached the young adult stage, then collected and washed three times using 50 mM NaCl. The worm pellet was resuspended in 1 ml of TRI Reagent (Sigma-Aldrich), vortexed for 15 min at room temperature, and stored at -80°C. RNA was isolated from TRIzol according to the manufacturer’s instructions. The samples were additionally purified with KAPA Pure magnetic beads (RNA to beads ratio: 1:3 v/v), and RNA quality was assessed using the Agilent TapeStation system. Samples for wild-type and *tent-5(tm3504)* males and hermaphrodites were prepared in two independent biological replicates.

### Direct RNA sequencing and data analysis

Sequencing libraries were prepared using the DRS Kit (Oxford Nanopore Technologies, SQK-RNA002) according to the manufacturer’s protocol. In each library, mRNA isolated from male or hermaphrodite worms (0, 5 - 1, 5 μg depending on the amount of isolated RNA) was mixed with 2 ng of *in vitro* transcribed poly(A) standards. Sequencing was performed with the MinION device, followed by basecalling with Guppy 6.0.0 (ONT), mapping to WBCel235 reference (MiniMap v2.17 (25) with options -k 14 -ax map-ont –secondary = no), and processing with samtools v1.9 (26). Lengths of poly(A) tails were estimated with Nanopolish (v0.13.2) as described before (13, 16). Poly(A) length distributions were compared between conditions for transcripts with a minimum of 5 reads per condition using the Wilcoxon test. Adjusted *p*-values were calculated using the Benjamini-Hochberg method. Differential expression analysis was performed using DESeq2 v1.28 Bioconductor package (27). GO enrichment analysis was performed using WormBase Enrichment Suite (28). The subcellular localization of proteins was assessed with DeepLoc v2.0 (29). DRS data have been deposited to the European Nucleotide Archive (ENA) with the following accession numbers: ERS20270802, ERS20271308, ERS20270803, ERS20271309, ERS20270804, ERS20270805, ERS20227048, and ERS20270807.

The dataset from Ebbing A. *et al.* (2018) for comparisons of sex-enriched genes was reanalyzed from raw *fastq* files deposited by authors in Gene Expression Omnibus (GEO). Reads were mapped to WBCel235 reference (STAR v2.7.10a (30)) and followed by processing with samtools v1.9 (26). Read counts were collected using featureCounts from the Subread package (v2.0.6 (31) with options -Q 10 -p -C -B and WBCel235 annotation). Differential expression analysis was performed using DESeq2 v1.28 Bioconductor package (27).

### Spermatids isolation

A day before the spermatids isolation, a few wild-type or *tent-5(tm3504)* L4 stage males were transferred to fresh NGM plates seeded with *E. coli* HB101. Separating males from hermaphrodites allows spermatids to accumulate in the male’s gonad before the experimental procedure (32). Males were dissected the next day in a drop of sperm medium (50 mM HEPES, 50 mM NaCl, 25 mM KCl, 5 mM CaCl_2_, 1 mM MgSO_4_, and 1 mg/ml BSA) by cutting the posterior end of the male with a needle. The isolated spermatids were visualized immediately (32, 33).

### Microscopic observations

Isolated spermatids were imaged using the Nikon Eclipse 80i microscope with a 100×/oil immersion lens. The spermatids’ size was assessed using ImageJ2 software (34) and compared between wild-type and *tent-5(tm3504)* males using a two-tailed unpaired Student’s *t*-test. To evaluate the localization of TENT-5 in males, *tent-5::gfp* worms were immobilized with 25 μM levamisole on freshly prepared 2% agarose pads and immediately imaged on the Zeiss LSM800 confocal microscope with a 40×/oil immersion lens.

### Fertility and mating behavior experiments

To assess male fertility, wild-type or *tent-5(tm3504)* L4 stage males were crossed with L4 stage *fog-2(oz40)* females (which are unable to produce their own sperm (35)). For each cross, one female and four males were placed on an NGM plate with a 5-mm diameter *E. coli* lawn. Crosses were performed at 20°C and continued throughout the entire female lifespan. The number of progeny was counted and compared between wild-type and *tent-5(tm3504)* males using a two-tailed unpaired Student’s *t*-test. For comparison of mating behavior, wild-type or *tent-5(tm3504)* young adult males were crossed with young adult *unc-45(m94)* hermaphrodites (uncoordinated hermaphrodites were used to increase mating efficiency (36)). For each cross, ten hermaphrodites and five males were placed on an NGM plate seeded with *E. coli* HB101 and recorded for 30 minutes. Recorded movies were subsequently analyzed, and mating behavior was described according to the following criteria: 1 – number of different hermaphrodites touched by the male, 2 – number of contacts with any hermaphrodite, 3 – backward locomotion after touching the hermaphrodite with the tail, 4 – turning around the hermaphrodite’s body, 5 – successful location of the hermaphrodite’s vulva, and 6 – successful spicule insertion (36, 37). The number of each incident was then compared between wild-type and *tent-5(tm3504)* males using a two-tailed unpaired Student’s *t*-test (with or without Welch’s correction depending on variance distributions).

## Results

### Male-enriched genes are expressed mainly in sperm, seminal vesicle, and vas deferens

It is widely recognized that gene expression displays significant variability depending on the sex of the organism, contributing to differences in appearance and behavior. Studies of sex-specific gene expression have been conducted in various organisms, including the nematode *C. elegans* (38–42). Here, we used Oxford Nanopore Direct RNA Sequencing (DRS) to expand on previous studies by simultaneously investigating differences in both gene expression and mRNA poly(A) tail length dynamics between young adult *C. elegans* wild-type males and hermaphrodites. Our DRS analysis revealed substantial differences in gene expression between the two sexes, with 1538 genes exhibiting sex-dependent expression patterns (Figure 1A, Supplementary Table S1). Among these, 1203 genes were enriched in males, while 335 were enriched in hermaphrodites. To validate our findings, we compared our results with previously published datasets that utilized different approaches to male collection and sample preparation. For instance, Kim B. et al. (2016) identified 1739 genes enriched in *him-5* mutant males using Illumina sequencing (42), of which 802 genes overlapped with our study (Figure 1B, Supplementary Table S1). Similarly, Ebbing A. et al. (2018) detected 3529 genes upregulated in wild-type males (41), with 1090 genes overlapping with our dataset (Figure 1B, Supplementary Table S1). The substantial overlap between male-enriched genes across different studies underscores the robustness of our approach and the reliability of our data for elucidating sex-specific gene regulation in *C. elegans*.

**Figure 1.**
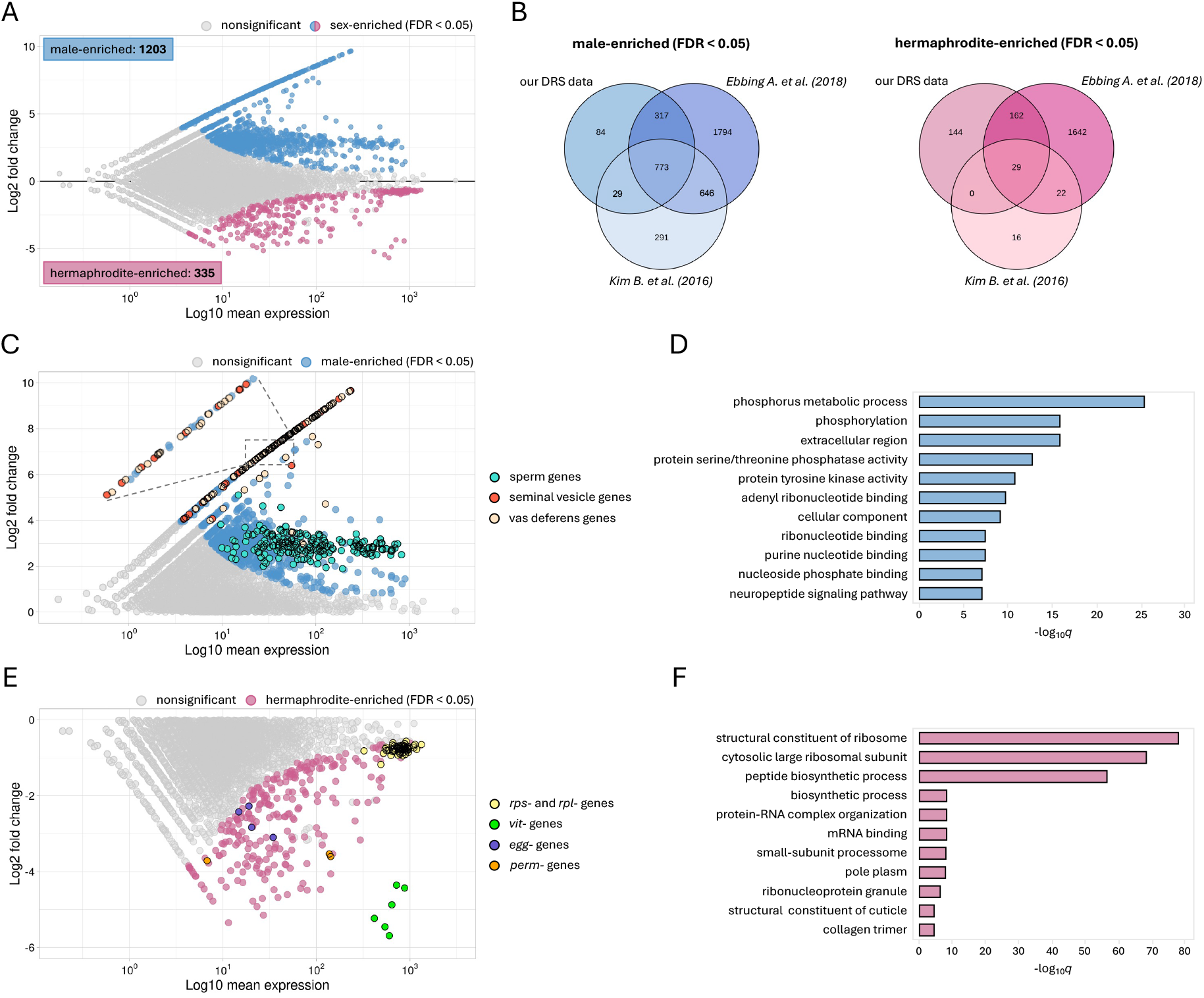
Differences in gene expression patterns between wild-type males and hermaphrodites. **(A)** MA plot illustrating differential gene expression between males and hermaphrodites of N2 wild-type worms. Significantly changed genes (FDR < 0.05) are marked with blue and pink dots for male-enriched and hermaphrodite-enriched genes, respectively. **(B)** Venn diagrams showing overlaps between our DRS data and two other studies comparing transcriptomes of *C. elegans* males and hermaphrodites (Kim B. *et al.* 2016 and Ebbing A. *et al.* 2018) (41, 42). For each compared group, the significantly enriched pool of genes was included according to the cut-off FDR value set in each study. The overlap for male-enriched genes is shown in blue and for hermaphrodite-enriched genes in pink. **(C)** Zoom on the half of the MA plot showing genes upregulated in males. Significantly changed genes (FDR < 0.05) are marked with blue dots. Among these, three of the most interesting groups of genes were selected: sperm genes (cyan dots), seminal vesicle genes (red dots), and vas deferens genes (beige dots). Groups of genes were defined based on Ebbing A. *et al.* 2018 (41). A zoomed-in fragment of the plot is shown for clearer visualization of overlapping dots. **(D)** Top GO terms for the male-enriched genes ordered by adjusted *p*-value (WormBase Enrichment Suite). **(E)** Zoom on the half of the MA plot showing genes upregulated in hermaphrodites. Significantly changed genes (FDR < 0.05) are marked with pink dots. Four groups of genes are marked: ribosomal genes (*rpl* and *rps*, yellow dots), vitellogenin genes (*vit*, green dots), *egg* genes (purple dots), and *perm* genes (orange dots). **(F)** Top GO terms for the hermaphrodite-enriched genes ordered by adjusted *p*-value (WormBase Enrichment Suite).

We then analyzed the gene sets enriched in males and hermaphrodites in more detail. The Gene Ontology (GO) term analysis of male-enriched genes revealed a distinctive association with phosphatase and kinase activities (Figure 1C and D), consistent with previous reports identifying these as characteristic features of sperm-related genes (42–45). Indeed, among male-enriched genes, we detected a large population encoding sperm components: 30 genes from the *msp* family, 17 from the *ssp* family, and 9 from the *nspd* family (Supplementary Table S1). While both *C. elegans* sexes generate sperm, males exhibit a prolonged sperm production period, resulting in higher overall sperm levels (46). This accounts for the observed upregulation of sperm genes in males compared to hermaphrodites (Figure 1C). Intriguingly, apart from the sperm genes, many other male-enriched genes are exclusively expressed in males and encode proteins with unknown functions, presenting an opportunity for further exploration in *C. elegans* male research. Subsequent analyses of genes specific to males allowed us to categorize them into two major groups: seminal vesicle genes and vas deferens genes (Figure 1C). The seminal vesicle and vas deferens are male-specific tissues that are responsible for producing seminal fluid and transporting it into the hermaphrodite uterus during mating (41). Despite the crucial role of these tissues in reproduction, the precise molecular regulatory mechanisms governing their functions are still not fully understood. However, it is known that due to their secretory function, male-specific tissues are rich in endoplasmic reticulum (ER) (47). In line with that, we observed that 82% of male-specific transcripts (defined as those with no expression in hermaphrodites, FDR < 0.05) are predicted to contain a signal peptide-encoding sequence targeting them to the secretory pathway through the ER (Supplementary Figure S1, Supplementary Table S1).

The pool of hermaphrodite-enriched genes and their overlap with datasets from other studies appeared relatively small compared to that of males (Figure 1A and B). However, the genes we identified as enriched closely align with hermaphrodite reproductive functions. As expected, among the genes highly enriched in hermaphrodites (Supplementary Table S1), we found many implicated in hermaphrodite reproduction, germline function, and early embryo development. Notably, all six members of the vitellogenin (*vit*) family exhibited high enrichment in hermaphrodites (Figure 1E). These genes are predominantly expressed in the intestine of young adult and adult hermaphrodites, encoding yolk proteins responsible for transporting essential nutrients to developing embryos within the gonad (48). Similarly, genes from the *egg* family (e.g., *egg-1*, *egg-2*, *egg-5*, and *egg-6*), *perm* family (e.g., *perm-2*, *perm-4*), and proteins involved in RNA regulation (e.g., *gld-1, lin-41, puf-5, puf-7, puf-11, mex-5, mex-6*), which are all known to influence germ cell proliferation, differentiation, and germline function in hermaphrodites (49), showed substantial enrichment compared to males. Interestingly, we also observed significant differences in the expression of genes encoding ribosomal proteins (*rpl* and *rps*) between the sexes (Figure 1E and F). Some of these genes (e.g., *rpl-10*, *rpl-3*, *rpl-7A*, *rpl-9, rpl-34, rps-8, rps-11, rps-12, rps-3*) were proposed to play critical roles in germline and embryonic development (49). More broadly, ribosomal proteins are essential for protein synthesis and cell growth. We suspect that their differential expression between sexes might reflect differences in cellular metabolism and growth between hermaphrodites and males.

In conclusion, our DRS results reveal profound differences in gene expression between *C. elegans* sexes, consistent with previous studies (41, 42). Our approach provides a more robust basis for examining sex-related transcriptomes by analyzing a large population of wild-type males, thus avoiding the potential influence of the *him* (high incidence of males) mutation background on gene expression and poly(A) tail distributions. Notably, we observed the most pronounced sex-dependent expression for genes encoding poorly studied secreted proteins localized in male-specific tissues.

### Poly(A) tail distributions differ for males and hermaphrodites

Given that the poly(A) tail length often correlates with a transcript’s stability and expression level (8), we hypothesized that significant differences would be observed between the two sexes of *C. elegans* not only in gene expression but also in mRNA poly(A) tail length distribution. Utilizing the same DRS-based dataset of mRNAs isolated from hermaphrodites and males as described above, we found global mRNA poly(A) tail length profiles consistent with previous studies (Supplementary Table S2) (11–13, 50). Additionally, we identified a slight difference between the sexes, with global median poly(A) tail lengths of 51 nt for hermaphrodites and 55 nt for males (Figure 2A). Further analysis of poly(A) tail length distributions for differentially polyadenylated mRNAs (at least 5 nt difference between sexes, FDR < 0.05) revealed 208 transcripts with significantly longer poly(A) tails in hermaphrodites and 155 transcripts with longer poly(A) tails in males (Figure 2B, Supplementary Figure S2A and B, Supplementary Table S2). Interestingly, in males, a significant portion of transcripts with longer tails are lowly expressed (Figure 2C). This group of mRNAs primarily contributes to the longer median poly(A) tail length observed in males (Figure 2D). In contrast, many transcripts with longer tails in hermaphrodites encode highly abundant proteins, such as ribosomal proteins, collagens, or actin (Figure 2C and E, Supplementary Figure S2C, Supplementary Table S2). These genes were also upregulated in hermaphrodites compared to males based on our differential expression analysis (Figure 1E and F, Supplementary Table S1). We speculate that the elongation of tails for these transcripts may be attributed to the increased demand for their respective proteins to support embryo development in the hermaphrodite’s gonad.

**Figure 2.**
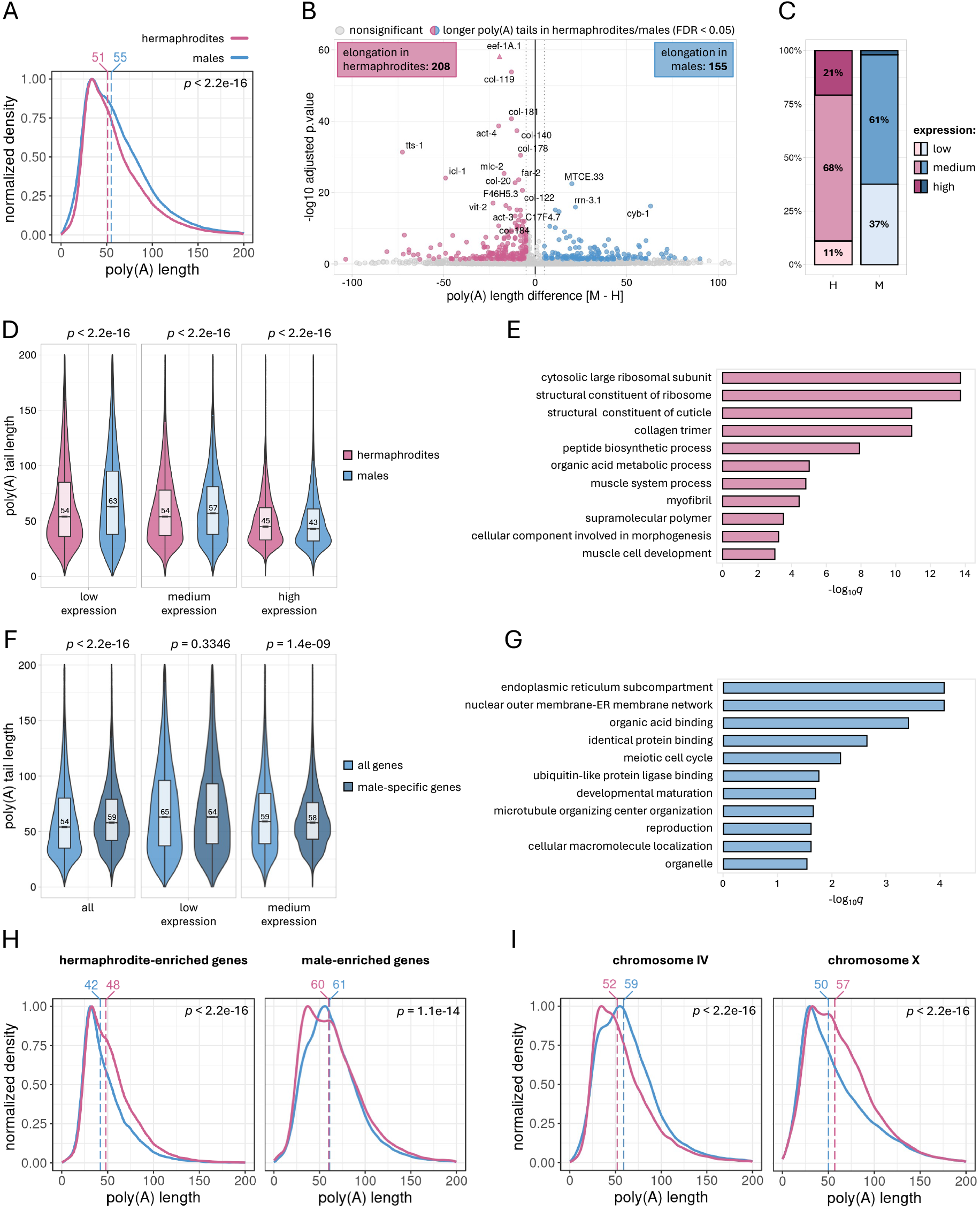
Differences in poly(A) distributions between wild-type males and hermaphrodites. **(A)** Density plot showing global differences in the poly(A) tail distribution between males (blue line) and hermaphrodites (pink line). Vertical dashed lines represent the median poly(A) tail length for each condition (in nucleotides). The plot was generated for all identified transcripts and normalized to 1. **(B)** Volcano plot displaying differential polyadenylation between males and hermaphrodites for N2 wild-type worms. Genes with significantly changed median poly(A) tail length (FDR < 0.05) are marked with blue dots (elongated in males) and pink dots (elongated in hermaphrodites). Triangles represent data points outside the *y-* axis limit. “M” in the *x*-axis title represents males and “H” represents hermaphrodites. **(C)** Fractions of genes with elongated poly(A) tails in hermaphrodites (pink bars) or males (blue bars), characterized by low, medium, or high expression levels. Expression levels are defined by baseMean values presented in Supplementary Table S1. Genes with baseMean < 20 are defined as lowly expressed, with 20 < baseMean < 500 as medium expressed, and with baseMean > 500 as highly expressed. “M” in the *x*-axis title represents males and “H” represents hermaphrodites. **(D)** Violin plots showing distributions of poly(A) tail length for genes grouped by their expression levels as defined in panel **C**. Numbers inside the boxes indicate the median poly(A) tail length for each condition. **(E)** Top GO terms for genes with poly(A) tails elongated in hermaphrodites ordered by adjusted *p*-value (WormBase Enrichment Suite). **(F)** Violin plots illustrating the distribution of poly(A) tail lengths for male-specific genes compared to other male-expressed genes. Male-specific genes are defined as those with no expression in hermaphrodites based on our DRS data (Supplementary Table S1). The left plot represents the distribution for all male-specific genes contrasted with all others, while the middle and right plots represent similar comparisons within groups of genes with similar expression levels as defined in panel **C**. No male-specific genes met our criteria for highly expressed genes. Numbers inside the boxes indicate the median poly(A) tail length for each condition. **(G)** Top GO terms for genes with poly(A) tails elongated in males ordered by adjusted *p*-value (WormBase Enrichment Suite). **(H)** Density plots showing differences in the poly(A) tail distribution for sex-enriched genes between males (blue line) and hermaphrodites (pink line). Vertical dashed lines represent the median poly(A) tail length for each condition (in nucleotides). Plots were generated for all transcripts significantly upregulated in either hermaphrodites or males based on Supplementary Table S1 and normalized to 1. **(I)** Density plots showing differences in the poly(A) tail distribution for all genes localized on chromosome IV or X between males (blue line) and hermaphrodites (pink line). Vertical dashed lines represent the median poly(A) tail length for each condition (in nucleotides). Plots were generated for all transcripts assigned to a particular chromosome and normalized to 1.

We observed that in males, poly(A) tail elongation is directed towards transcripts functionally associated with the endoplasmic reticulum (Figure 2G), suggesting the importance of efficient secretory machinery for proper male physiology. Since most male-specific genes encode proteins secreted by the ER (Supplementary Figure S1A, Supplementary Table S1), we analyzed their poly(A) tails. We found that globally, male-specific mRNAs exhibit significantly longer poly(A) tails compared to other transcripts expressed in males (Figure 2F, left panel). This observation can be explained by the much higher proportion of lowly expressed genes in this group, which generally have longer poly(A) tails than highly expressed ones (Figure 2D). Further comparisons of poly(A) tails within groups of genes with similar expression levels revealed no significant difference in median tail length between male-specific and other male-expressed genes (Figure 2F, middle and right panels). To investigate further, we also explored the relationship between poly(A) tail lengths and mRNA expression levels. For both hermaphrodites and males, we observed a negative correlation between mRNA tail length and expression level (Figure 2D and F), which is consistent with previous reports (11). Moreover, we found that transcripts significantly upregulated in hermaphrodites also carry globally longer poly(A) tails in hermaphrodites than in males (Figure 2H). A similar trend was observed for male-enriched transcripts (Figure 2H), which proves the importance of poly(A) tail in regulating gene expression levels.

Finally, due to the distinct chromosome compositions of males and hermaphrodites, we examined the global poly(A) length distributions based on gene location within each chromosome. Surprisingly, we identified differences in poly(A) profiles for transcripts from chromosomes IV and X (Figure 2I, Supplementary Figure S2D). Specifically, mRNAs from chromosome IV exhibited significantly longer poly(A) tails in males compared to hermaphrodites, whereas transcripts from chromosome X showed the opposite effect. Although we compared multiple features of genes encoded on each chromosome (expression levels, presence of sequence encoding signal peptides, GC content, transcript length, or 5′ UTR and 3′ UTR length) (Supplementary Figure S3), we were unable to explain how observed sex-dependent polyadenylation is influenced by gene chromosome location.

In addition to the significant differences in gene expression between males and hermaphrodites, our DRS results unveil distinct patterns of poly(A) tail length distribution between the two sexes. These findings underscore the important role of poly(A) metabolism in sex-dependent physiological processes.

### TENT-5 regulates the expression of male-specific genes

Within the cell, poly(A) tails can undergo modification by various proteins that either lengthen or shorten them (4–6). We hypothesized that the observed differences in poly(A) distributions between *C. elegans* males and hermaphrodites might result from the sex-specific activities of these proteins. To identify a potential candidate protein, we analyzed our DRS data to determine if the expression of known poly(A) polymerases and deadenylases varied between the sexes. We observed no significant differences in gene expression levels for any of these poly(A)-modifying enzymes (Figure 3A, Supplementary Table S1).

**Figure 3.**
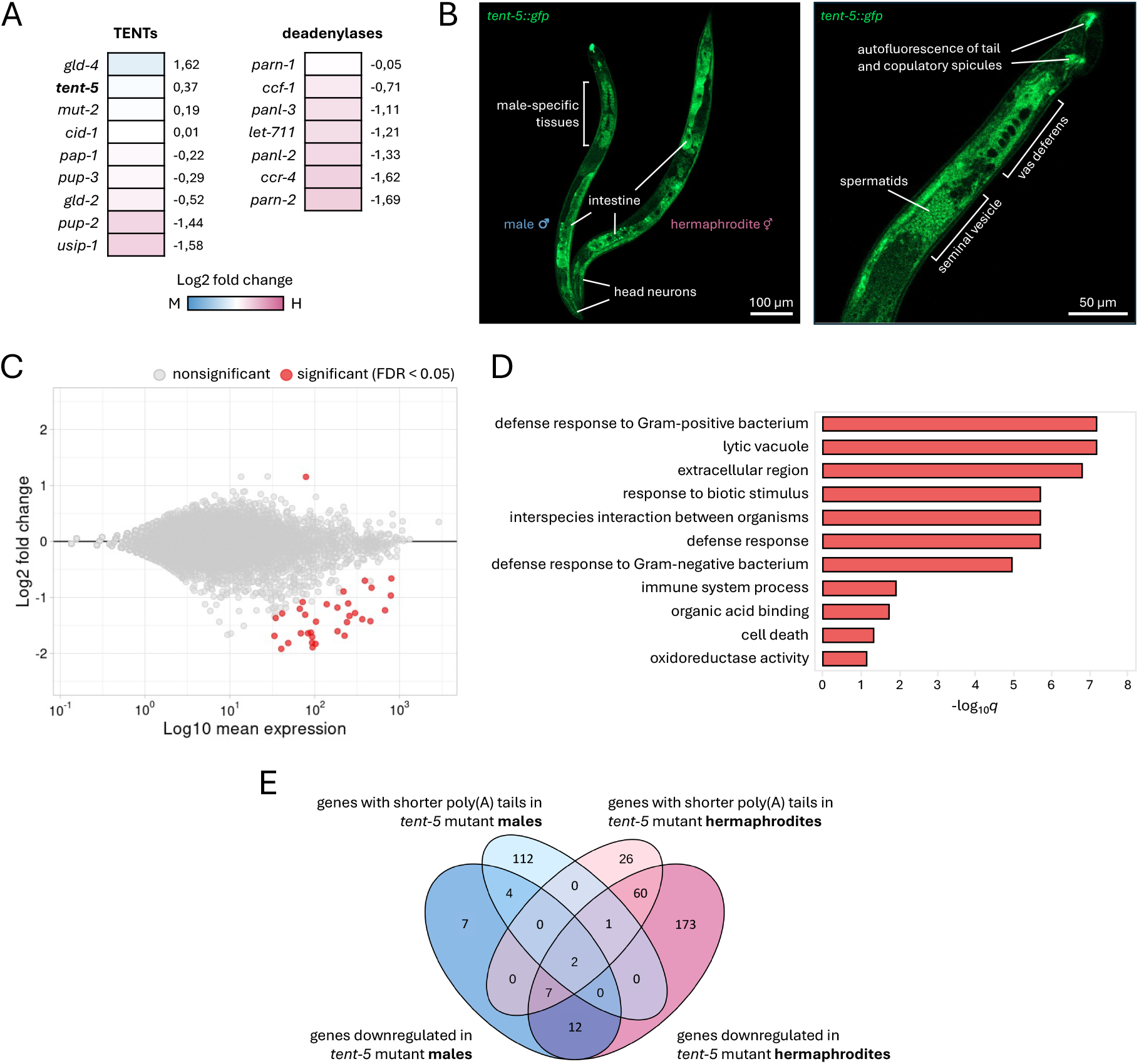
Differences in gene expression patterns between *tent-5* mutant and wild-type males. **(A)** Heatmap showing expression levels of TENTs and deadenylases between males and hermaphrodites. The color of the heatmap specifies whether a gene is upregulated in males (blue) or hermaphrodites (pink). The numbers on the right represent the log2 fold change value derived from Supplementary Table S1. “M” in the heatmap legend represents males and “H” represents hermaphrodites. **(B)** Fluorescence microscopic images of TENT-5-GFP expression in hermaphrodites and males (*tent-5(rtt6[tent-5::gfp::3xflag]* I). The left image represents differences and similarities in TENT-5 expression between sexes. It is composed of six separate shots to show the full body of a male and hermaphrodite. The right image shows the tail-end of a male body. Images were taken with 40x lens. **(C)** MA plot illustrating differential gene expression between wild-type and *tent-5* mutant worms. Significantly changed genes (FDR < 0.05) are marked with red dots. **(D)** Top GO terms for genes significantly downregulated in *tent-5* mutant males ordered by adjusted *p*-value (WormBase Enrichment Suite). **(E)** Venn diagram showing overlaps between our DRS data and our previous study on *tent-5* mutant hermaphrodites (Liudkovska V. *et al.* 2022) (13). Four different conditions were compared: genes downregulated in *tent-5* mutant males (blue), genes with shorter poly(A) tails in *tent-5* mutant males (light blue), genes downregulated in *tent-5* mutant hermaphrodites (pink), and genes with shorter poly(A) tails in *tent-5* mutant hermaphrodites (light pink). Included were all significantly downregulated genes (FDR < 0.05) (for differential expression) and all transcripts with significantly shorter poly(A) tail by a minimum of 5 nucleotides (FDR < 0.05) (for differential polyadenylation).

Our previous work in hermaphrodite *C. elegans* demonstrated that the polyadenylation of mRNAs encoding ER-targeted secreted proteins is mediated by the noncanonical poly(A) polymerase TENT-5 (13). Depletion of TENT-5 in hermaphrodites resulted in poly(A) tail shortening and downregulation of mRNAs encoding secreted innate immunity effector proteins, leading to increased susceptibility of *tent-5* mutants to pathogen infection (13). As mentioned above, our functional enrichment analysis showed that transcripts with longer poly(A) tails in males compared to hermaphrodites are associated with ER functionality (Figure 2F). Therefore, we assumed that TENT-5 might be responsible for their polyadenylation and thus contribute to the differential polyadenylation observed between the sexes, particularly evident for lowly expressed genes. Additionally, we hypothesized that since most male-specific transcripts are predicted to encode secreted proteins, TENT-5 might also regulate their expression, which could have been overlooked in our previous studies on hermaphrodites.

Using the *tent-5::gfp* knock-in strain, we examined TENT-5 localization in males. Confocal microscopy revealed a strong expression of TENT-5-GFP in the male intestine and head neurons (Figure 3B), consistent with previous observations in hermaphrodites (13). Notably, we detected a pronounced localization of TENT-5 in spermatids, seminal vesicles, and vas deferens (Figure 3B, Supplementary Figure S4A), where most of the male-enriched genes are expressed (Figure 1C). This observation suggested that, indeed, TENT-5 may regulate male-specific transcripts through their polyadenylation. Of note, our microscopic observations revealed intense autofluorescence at the male tale tip and copulatory spicules (Figure 3B, Supplementary Figure S4B), an intriguing feature previously briefly mentioned in the literature (51–53).

To identify TENT-5 substrates in *C. elegans* males and validate the hypothesis that TENT-5 regulates male-specific mRNAs and other transcripts encoding secreted proteins, we performed DRS on *tent-5* mutant and wild-type males, analyzing gene expression and poly(A) patterns. We observed significant dysregulation of 33 genes upon TENT-5 deletion in males: the expression levels of 32 genes were downregulated, and one was upregulated (Figure 3C, Supplementary Table S3). Among the downregulated genes, 21 were consistent with our previous findings in *tent-5*-deficient hermaphrodites (Figure 3E, Supplementary Table S3) (13), showing similar functional enrichment in defense response processes (Figure 3D). This underscores TENT-5’s role in *C. elegans* innate immunity, independently of nematode sex. However, poly(A) profiling of *tent-5* mutant males revealed more pronounced sex-related differences. Despite no global change in the poly(A) distribution between wild-type and *tent-5* mutant males (similar to our previous observations for hermaphrodites (13)) (Figure 4A), we detected 137 transcripts with altered poly(A) tail length (changed by at least 5 nt, FDR < 0.05), of which 119 had shorter and 18 longer poly(A) tails in *tent-5* mutant males (Figure 4B, Supplementary Figure S4C, Supplementary Table S4). Strikingly, among mRNAs with shortened poly(A) tails in mutants, 100 were male-specific, and 12 were male-enriched, accounting for 94% of mRNAs with shortened poly(A) tails in *tent-5* mutant males (Figure 4B, Supplementary Table S4). As expected, the majority of these transcripts are predicted to encode extracellular proteins (93%), a hallmark of TENT-5 substrates (13, 15, 16, 54, 55), which is even more pronounced in males than in hermaphrodites (Figure 4C and D, Supplementary Table S4).

**Figure 4.**
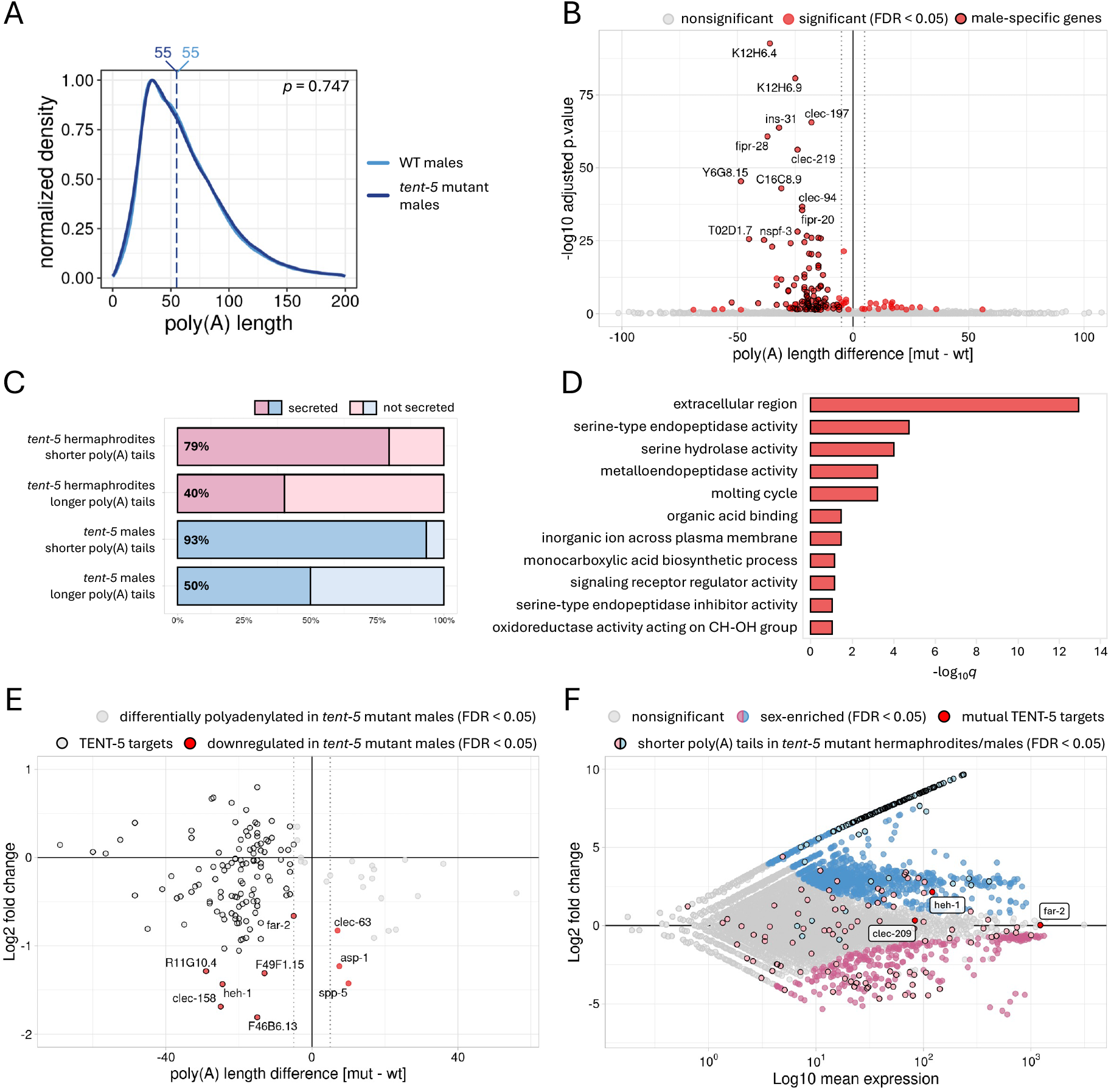
TENT-5 polyadenylates male-specific mRNAs encoding secreted proteins. **(A)** Density plot showing global differences in the poly(A) tail distribution between wild-type (blue) and *tent-5* mutant males (dark blue). Vertical dashed lines represent the median poly(A) tail length for each condition (in nucleotides). The plot was generated for all identified transcripts and normalized to 1. **(B)** Volcano plot showing differential polyadenylation between wild-type and *tent-5* mutant males. Transcripts with significantly changed median poly(A) tail length (FDR < 0.05) are marked with red dots. Black borderline is added to male-specific genes. Male-specific genes were all genes that had no expression in hermaphrodites based on our DRS data (Supplementary Table S1). **(C)** Fractions of genes with predicted extracellular localization by DeepLoc 2.0 software (29). For each condition, a group of significantly shortened/elongated transcripts by a minimum of 5 nucleotides was included. Data for hermaphrodites comes from Liudkovska V. *et al.* 2022 (13). **(D)** Top GO terms for genes with significantly shorter poly(A) tails in *tent-5* mutant males (by a minimum of 5 nucleotides) ordered by adjusted *p*-value (WormBase Enrichment Suite). **(E)** The relationship between the difference in the expression level and the difference in the median poly(A) tail length of respective transcripts for *tent-5* mutant compared to wild-type males. TENT-5 targets (genes with significantly shorter poly(A) tail in *tent-5* mutant worms by a minimum of 5 nucleotides; FDR < 0.05) are highlighted with black borderline. Genes significantly downregulated in *tent-5* mutant males (FDR < 0.05) are marked with red dots. **(F)** MA plot showing differential gene expression between males and hermaphrodites of N2 wild-type worms (Figure 1A). Significantly changed genes (FDR < 0.05) are marked with blue and pink dots for male-enriched and hermaphrodite-enriched genes, respectively. Additionally, marked are TENT-5 targets (genes with significantly shorter poly(A) tail in *tent-5* mutant worms by a minimum of 5 nucleotides; FDR < 0.05) in males (light blue, black borderline) and hermaphrodites (light pink, black borderline; based on Liudkovska V. *et al.* 2022). Red dots with labels represent common TENT-5 targets for both sexes.

Approximately 66% of TENT-5 targets encode putative components of seminal fluid expressed in male-specific tissues (Supplementary Table S4), suggesting the importance of TENT-5-mediated polyadenylation in regulating certain aspects of male reproduction. For instance, some of these substrates possess serine-type endopeptidase activity (Figure 4D), which has been previously linked with sperm activation in *C. elegans* (33, 56). Although the process of sperm activation is not well understood in nematodes, it is presumed that inactive spermatids stored in seminal vesicle are transported through the male’s vas deferens and cloaca into the hermaphrodite’s uterus. After ejaculation, spermatids are activated by seminal fluid proteins to form functional sperm (57). So far, only a few genes involved in this process have been reported (33, 56). Interestingly, one of them (*try-5*) is polyadenylated by TENT-5 based on our results (Supplementary Figure S4D, Supplementary Table S4).

Among the mRNAs with shortened poly(A) tails in *tent-5* mutant males, we also detected 22 members of the *clec* family (C-type lectins) (Supplementary Figure S4D, Supplementary Table S4), which are reported to be immune effectors (58, 59). The enrichment of C-type lectins in the seminal fluid suggests its possible function in the nematode’s defense response. Recent studies have shown that *C. elegans* males are more resistant to fungal, viral, and bacterial infections, as well as to heat, osmotic, and oxidative stress (60). Also in other species, seminal fluid has been shown to have antimicrobial properties, helping to protect both female and male reproductive tracks against infections (61, 62). This raises the interesting hypothesis that *C. elegans* males utilize seminal fluid genes for both defense response and mating purposes (57). However, the exact mechanism of that possible parallel regulation remains to be established.

Additionally, we analyzed how differential gene expression correlates with poly(A) tail length. Although only six genes displayed both significant downregulation and shorter poly(A) tails in mutant males (Figures 3E and 4E), we observed a trend where TENT-5-mediated poly(A) tail elongation tends to stabilize the corresponding mRNA (Figure 4E), a pattern also noted in hermaphrodites (13). This finding suggests that the correlation between mRNA poly(A) tail length and expression level might be largely tissue-specific or specific to certain transcript types or groups. Surprisingly, the overlap between transcripts with shortened poly(A) tails in *tent-5* mutant males and hermaphrodites was minimal (Figures 3E and 4F), with only three genes (*heh-1*, *far-2,* and *clec-209*) emerging as shared TENT-5 targets across sexes (Supplementary Figure S4D, Supplementary Table S4). Moreover, *heh-1* and *far-2* were also significantly downregulated in both sexes (Figure 3E). Although not well studied, these genes are predicted to encode proteins responsible for sterol or fatty acid binding (49). Interestingly, male-specific tissues, where TENT-5 activity in males is most pronounced, are known to be cholesterol-rich (63). Therefore, we speculate that TENT-5 might be involved in cholesterol metabolism in both sexes by stabilizing *heh-1* and *far-2* mRNAs.

Overall, TENT-5 activity in males focuses almost exclusively on polyadenylating male-specific transcripts, whereas in hermaphrodites, it appears much more versatile (Figure 4F). The functions of most male-specific TENT-5 substrates remain unknown, making it difficult to speculate about TENT-5’s physiological role in males. However, we expect that the elongation of poly(A) tails of so many putative seminal fluid components in males is crucial for their reproduction.

### TENT-5-mediated poly(A) tail elongation does not affect male mating behavior

Following our observation that TENT-5 is responsible for the polyadenylation of male-specific transcripts in *C. elegans* males, we attempted to identify the resulting physiological phenotype. We hypothesized that since TENT-5 targets comprise presumed components of the seminal fluid, the phenotype of *tent-5* mutants might exhibit abnormal mating phenotypes. Firstly, we monitored the mating behaviors of wild-type and *tent-5* mutant males. We did not detect any significant differences in the frequency of male contact with hermaphrodites, characteristic backward movements, vulva searching, or spicule insertion (Figure 5A, Supplementary Movie S1). The significant difference between wild-type and mutant males was noticed only for the turning behavior. However, we cannot exclude that this observation results from a random variance between individual worm behavior rather than from the TENT-5 depletion. The absence of the obvious mating behavioral phenotype indicates that *tent-5* mutant males have no impaired neuronal or pheromonal signaling. Next, we assumed that post-transcriptional changes in male-specific transcripts might influence the potency of the seminal fluid in properly activating male sperm post-ejaculation. To investigate this, we assessed male fertility by counting the offspring produced by *fog-2* females crossed with either wild-type or *tent-5* mutant males. Progeny numbers were comparable between the two conditions (Figure 5B), which suggests that TENT-5 does not significantly influence male mating behavior or efficiency.

**Figure 5.**
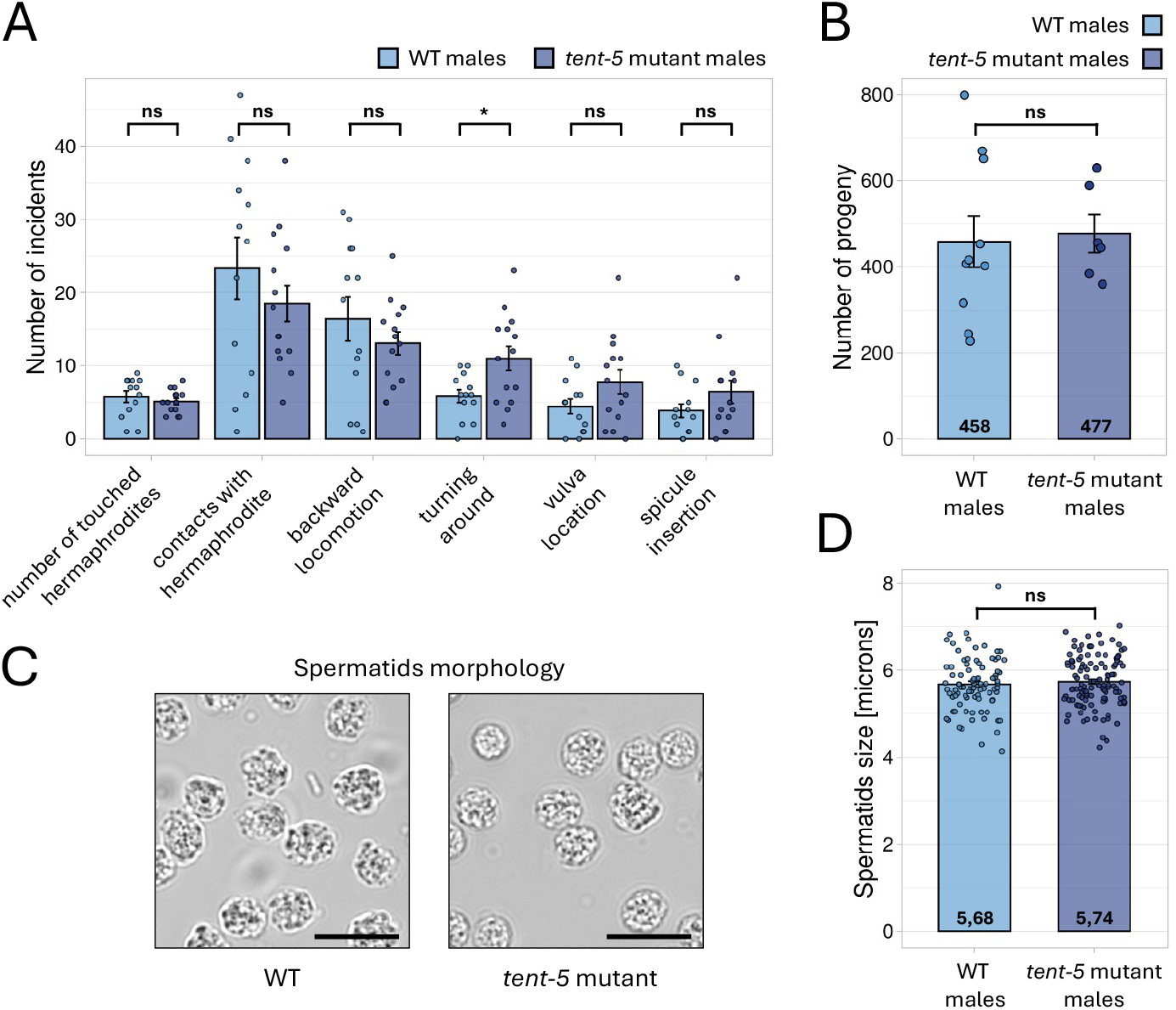
Comparison of mating behavior, fertility, and spermatids morphology between *tent-5* mutant and wild-type males. **(A)** Differences between the number of mate-related incidents performed by wild-type (n=15) and *tent-5* mutant (n=14) males. Barplots represent mean values with SD. ns => not significant; * => *p*-value < 0.05 (two-tailed *t-* test). **(B)** Differences in brood sizes between wild-type (n=10) and *tent-5* mutant (n=6) males crossed with *fog-2* females. Barplots represent mean values with SD. ns => not significant (two-tailed *t-*test). **(C)** Microscopic images of spermatids morphology for wild-type and *tent-5* mutant males. Images were taken with 100x magnification. Scale bars => 10 microns. **(D)** Differences in spermatids diameter between wild-type (n=86) and *tent-5* mutant (n=114) males. Barplots represent mean values with SD. ns => not significant (two-tailed *t-*test).

Knowing that some seminal fluid components are involved in the process of sperm activation (33, 56), we hypothesized that their TENT-5-mediated regulation might influence the morphology of spermatids stored in the male’s seminal vesicle. However, our microscopic observations did not show any alteration in spermatids morphology (Figure 5C, Supplementary Figure S5) and size (Figure 5D). We also did not detect any incidents of matured sperm, that were activated before the ejaculation. Nevertheless, considering the strong modifications in seminal fluid components observed at the transcriptome level, we would expect that these changes should manifest as some physiological phenotype, that were not examined in this study. More complex *in vivo* studies would need to be performed to elucidate which physiological processes are regulated by TENT-5 poly(A) polymerase in males. Unfortunately, as mentioned above, most of the predicted TENT-5 targets in males do not have any functional annotation, which makes it harder to anticipate their possible function in male physiology.

## Discussion

Advancements in methods that allow precise genome-wide estimation of the poly(A) tail length and composition have significantly boosted our understanding of poly(A) tail metabolism and its role in gene regulation (8). However, it quickly became evident that this regulation is highly cell- and tissue-type specific, and extremely dynamic in response to environmental and developmental cues. Therefore, it should be explored in comprehensive ways that can capture these dynamic changes. Several excellent studies leveraged these novel methodologies to profile poly(A) tail length during oocyte maturation in *Drosophila melanogaster* (18) and *Xenopus laevis* (64), mammalian oocyte-to-embryo transition (65), and embryonic development of *Danio regio*, *X. laevis*, and *C. elegans* (10, 12, 50). However, sex-related differences in global poly(A) metabolism remain largely overlooked in most organisms. In this study, we employed Direct RNA Sequencing to investigate poly(A) tail distribution differences between hermaphrodite and male *C. elegans*, aiming to understand their underlying reasons and consequences.

We observed clear, previously reported differences in gene expression between males and hermaphrodites, reflecting the distinct cellular metabolism, growth, and physiology of adult hermaphrodites and males. Additionally, our poly(A) profiling revealed a distinct sex-dependent distribution of poly(A) tail length. In general, our data align with previous observations that highly abundant housekeeping genes possess relatively short poly(A) tails (11, 19). We confirmed this negative correlation between mRNA poly(A) tail length and expression level for *C. elegans* males and hermaphrodites. Notably, highly expressed mRNAs have slightly shorter tails in males compared to hermaphrodites, whereas lowly expressed mRNAs have significantly longer tails in males. Interestingly, sex-enriched mRNAs (those expressed in both sexes but upregulated in one) tend to have longer tails in the respective sex. The mechanism behind this sex-dependent difference remains unclear. We also noted chromosomal differences in poly(A) distributions. Transcripts encoded by genes on the X chromosome have longer poly(A) tails in hermaphrodites, while those derived from chromosome IV have longer tails in males. The X chromosome is enriched with female transcripts and largely devoid of male-enriched ones (43, 66), likely leading to the observed global elongation of poly(A) tails for X chromosome transcripts in hermaphrodites. Conversely, the opposite trend for chromosome IV can be explained by the presence of many male-enriched, spermatogenesis-related genes (67). Another notable observation concerns male-specific transcripts. We found that 82% of these transcripts are predicted to contain signal peptide-encoding sequences targeting them to the secretory pathway through the ER. Interestingly, most of these transcripts have medium to low expression levels. When compared with transcripts of the same expression levels, male-specific mRNAs show no difference in their median poly(A) tail length. However, compared to all mRNAs expressed in males, male-specific transcripts tend to have longer poly(A) tails.

The mechanistic reasons for the sex-specific differences in poly(A) tail distributions are not yet fully understood. Multiple factors influence mRNA poly(A) tail length, with the enzymatic activities of deadenylases, noncanonical poly(A) polymerases, and terminal uridylyltransferases playing primary roles (4–6). The steady-state poly(A) distribution at the transcriptome scale is achieved through the cell- or tissue-specific concentrations of these enzymes and their varying affinities modulated by auxiliary factors toward particular transcripts. We discovered that for a specific group of mRNAs, which predominantly encode secreted components of seminal fluid, the poly(A) tail length is affected by the TENT-5 ncPAP. TENT-5 localizes to male reproductive tissues rich in ER, such as spermatids, seminal vesicle, and vas deferens, where it potentially stabilizes male-specific mRNAs. In hermaphrodites, the same enzyme is mainly expressed in the intestine and enhances the expression of genes encoding secreted innate immune effectors (13). Therefore, in both *C. elegans* sexes, TENT-5 polyadenylates transcripts encoding proteins with predicted extracellular localization, underscoring the notion that the TENT5 family of ncPAPs is essential for post-transcriptional regulation of mRNAs encoding proteins directed towards the ER secretory pathway (13, 15, 16, 54). As described in our previous work (13), the exact mechanism behind TENT-5-substrate specificity is not yet clear. However, as a fraction of TENT-5 resides in the ER, we proposed that TENT-5 specificity is directly or indirectly driven by its colocalization with ER-targeted mRNAs (13). We have previously predicted that cytoplasmic polyadenylation by TENT-5 would be most pronounced in tissues with high secretion capacity. Consequently, in males, it is not surprising that TENT-5 focuses on regulating male-specific genes expressed primarily in the ER-rich seminal vesicle and vas deferens. Furthermore, our data strongly suggests that TENT-5 affects the composition of seminal fluid by modulating its transcriptome, potentially playing a role in male reproduction. However, despite intensive efforts, we have not identified phenotypes related to fertility or mating behavior in TENT-5-deficient males, possibly because these phenotypes manifest only under specific environmental conditions. We also cannot exclude the possibility that other ncPAPs may redundantly function with TENT-5 specifically in male tissues. One potential candidate is GLD-4, which was slightly enriched in males according to our DRS (Figure 3A, Supplementary Table S1). It has been shown that while GLD-4 alone partially reduces the fertility of hermaphrodites, it does not affect spermatogenesis and male fertility (21). In the future, it would be interesting to analyze the poly(A) distribution in males devoid of both TENT-5 and GLD-4.

From an evolutionary perspective, it is interesting to examine the role of noncanonical cytoplasmic poly(A) polymerases in reproduction and gametogenesis across different animal species. In *C. elegans*, GLD-2 is a crucial regulator of the germline mitosis-to-meiosis transition and subsequent gametogenesis (20). Consequently, GLD-2 depletion in nematodes results in defective oocytes and sperm production, leading to infertility in both hermaphrodites and males (22). Similarly, two homologs of GLD-2 in *D. melanogaster*, Gld2 and Wispy, are essential for proper gamete production in a sex-dependent manner (23, 24). Gld2 is expressed exclusively in the male germline and is required for the completion of spermatogenesis; its absence results in complete sterility in male flies (23). Wispy, on the other hand, regulates late oogenesis and is crucial for female fly fertility (24). Surprisingly, the knockout of the mouse homolog of GLD-2, TENT2, does not cause abnormalities in gamete production or affect reproduction (68). In mammals, the TENT5 family of ncPAPs, which includes four members (TENT5A-D), plays an essential role in spermatogenesis and oogenesis (15). In mice, TENT5C and TENT5D are expressed in spermatocytes and spermatids, regulating different aspects of spermiogenesis (15, 69, 70). Depletion of either TENT5C or TENT5D results in complete male infertility (15). Additionally, TENT5C and TENT5B regulate oogenesis, and the double knockout mutation of both proteins leads to infertility (15). In contrast, TENT-5 poly(A) polymerase in worms is not essential for hermaphrodite fertility (13) and only slightly impacts the male reproductive system. These findings indicate that the role of cytoplasmic polyadenylation is highly conserved across species. However, different species may rely on the activities of different TENTs to varying extents.

Further research is needed to uncover the complexity of poly(A) tail metabolism at the organismal level, not only in gametogenesis and reproduction but also in many other physiological processes. Our DRS analysis represents an important step toward expanding the current understanding of cytoplasmic polyadenylation. Additionally, we believe our results provide a foundation for further studies on *C. elegans* male physiology.

## Data availability

The Nanopore Direct RNA Sequencing data have been deposited to the European Nucleotide Archive (ENA) with the following accession numbers: ERS20270802, ERS20271308, ERS20270803, ERS20271309, ERS20270804, ERS20270805, ERS20227048, and ERS20270807.

## Acknowledgments

We are grateful to the members of the Dziembowski group for their support and insightful comments. We thank Paweł Krawczyk and Natalia Gumińska for the assistance with bioinformatic analyses, Aleksandra Brouze and Seweryn Mroczek for running Nanopore sequencing on a subset of samples, Bartosz Tarkowski for discussing ideas, Wojciech Pokrzywa and Anwesha Sarkar for sharing *unc-45(m94)* strain, and Tomasz Węgierski for the support with microscopic observations.

## Funding

This work was supported by the National Science Center (OPUS 17 UMO-2019/33/B/NZ2/01773) and Horizon Europa (European Research (AdG nr 101097317). Some *C. elegans* strains were provided by the CGC, which is funded by NIH Office of Research Infrastructure Programs (P40 OD010440).

## Author contributions

A.D. provided funding and directed the project. Z.M, V.L, and A.D designed the experiments. Z.M conducted all *C. elegans* experiments and analyzed the sequencing data. Z.M drafted the initial manuscript. V.L and A.D revised and edited the manuscript.

## Notes

### Competing Interest Statement

The authors have declared no competing interest.

