## Supplementary Figures for "TENT-5 regulates the expression of male-specific genes in *Caenorhabditis elegans*"

### Supplementary Materials

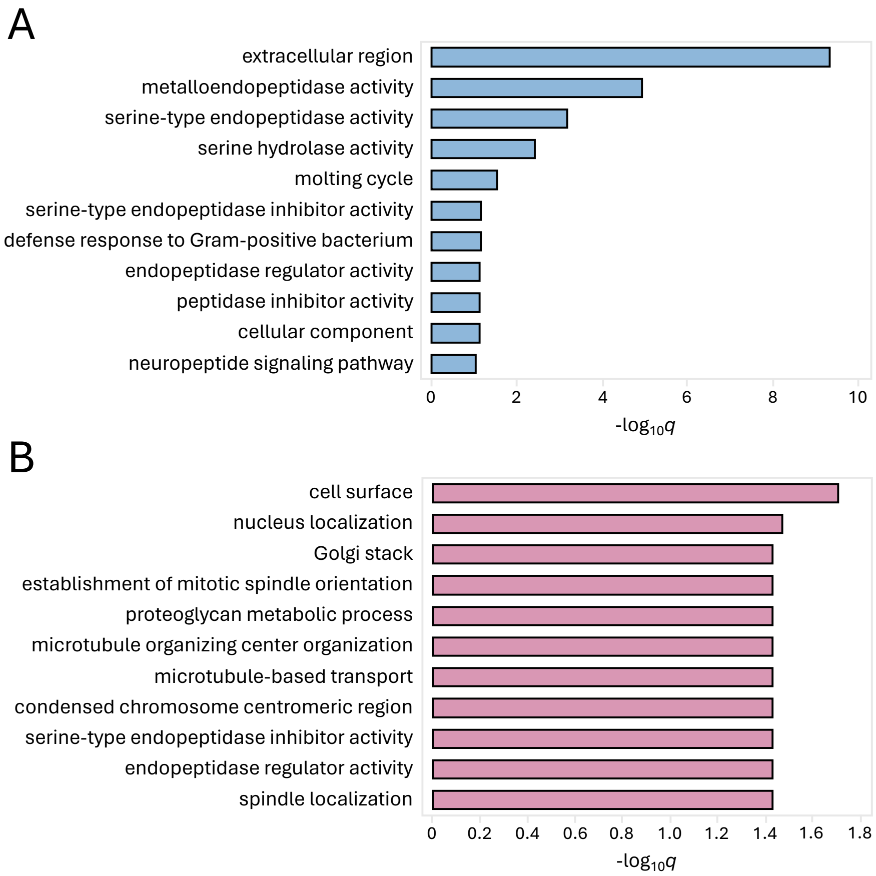


**Supplementary Figure S1. Functional enrichment for groups of male- and hermaphrodite-specific genes.**
**(A)** Top GO terms for male-specific genes with FDR < 0.05 ordered by adjusted *p*-value (WormBase Enrichment Suite).
**(B)** Top GO terms for hermaphrodite-specific genes with FDR < 0.05 ordered by adjusted *p*-value (WormBase Enrichment Suite).

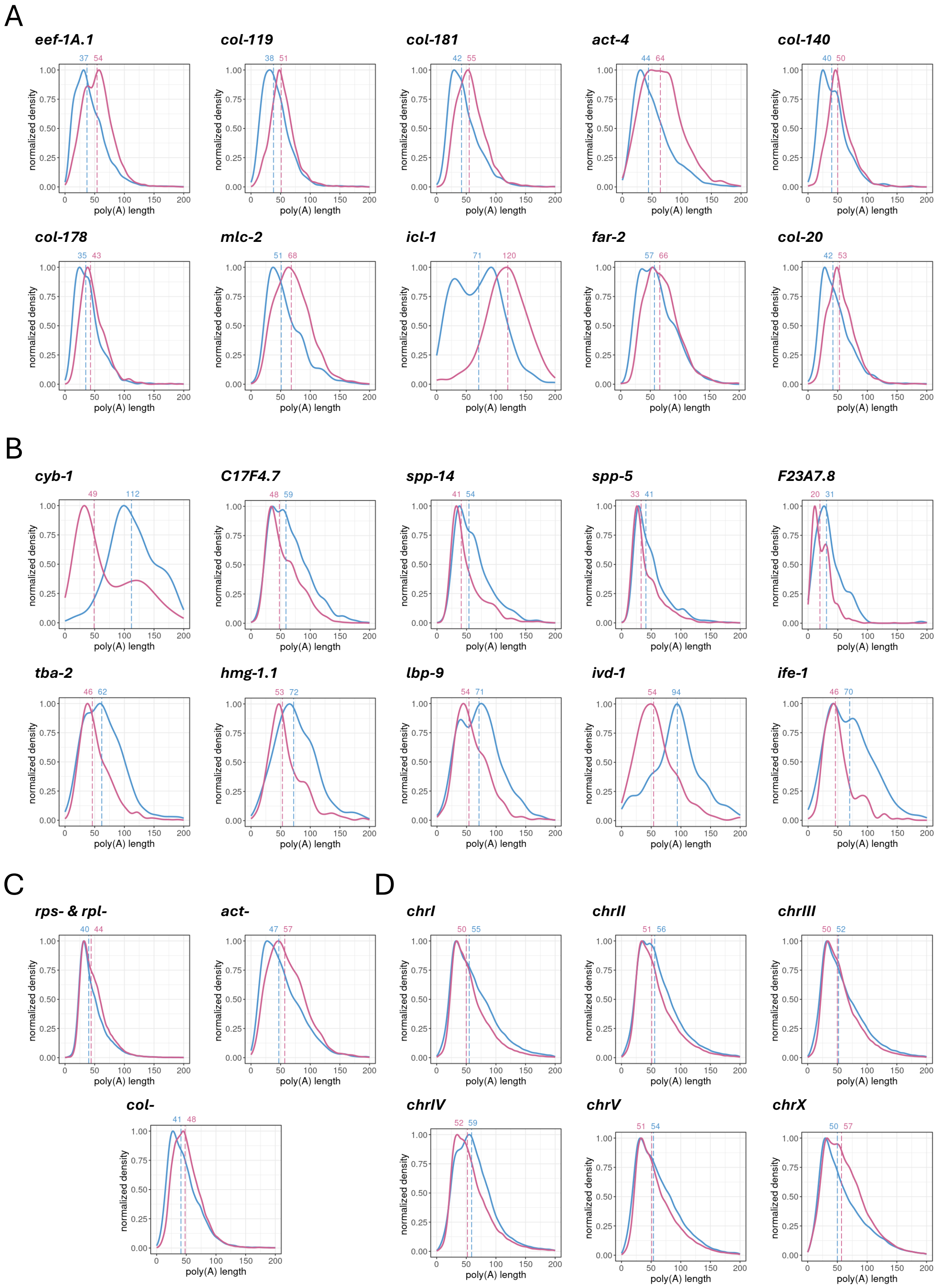


**Supplementary Figure S2. Differences in poly(A) distributions between wild-type males and hermaphrodites.**

**(A)** Top ten protein-coding transcripts with the most significant poly(A) elongation in hermaphrodites according to adjusted *p*-value (Supplementary Table S2). Poly(A) distributions are shown as blue and pink lines for males and hermaphrodites, respectively. Vertical dashed lines represent the median poly(A) tail length for each condition (in nucleotides). The plots were normalized to 1.

**(B)** Top ten protein-coding transcripts with the most significant poly(A) elongation in males according to adjusted *p*-value (Supplementary Table S2). Poly(A) distributions are shown as blue and pink lines for males and hermaphrodites, respectively. Vertical dashed lines represent the median poly(A) tail length for each condition (in nucleotides). The plots were normalized to 1.

**(C)** Density plots showing differences in the global poly(A) tail distribution for ribosomal genes (*rpl* and *rps*), actin genes (*act*), and collagen genes (*col*) between males (blue line) and hermaphrodites (pink line). Vertical dashed lines represent the median poly(A) tail length for each condition (in nucleotides). The plots were generated for all transcripts identified from each group and normalized to 1.

**(D)** Density plots showing differences in the global poly(A) tail distribution for genes localized on all six chromosomes between males (blue line) and hermaphrodites (pink line). Vertical dashed lines represent the median poly(A) tail length for each condition (in nucleotides). Plots were generated for all transcripts assigned to a particular chromosome and normalized to 1.

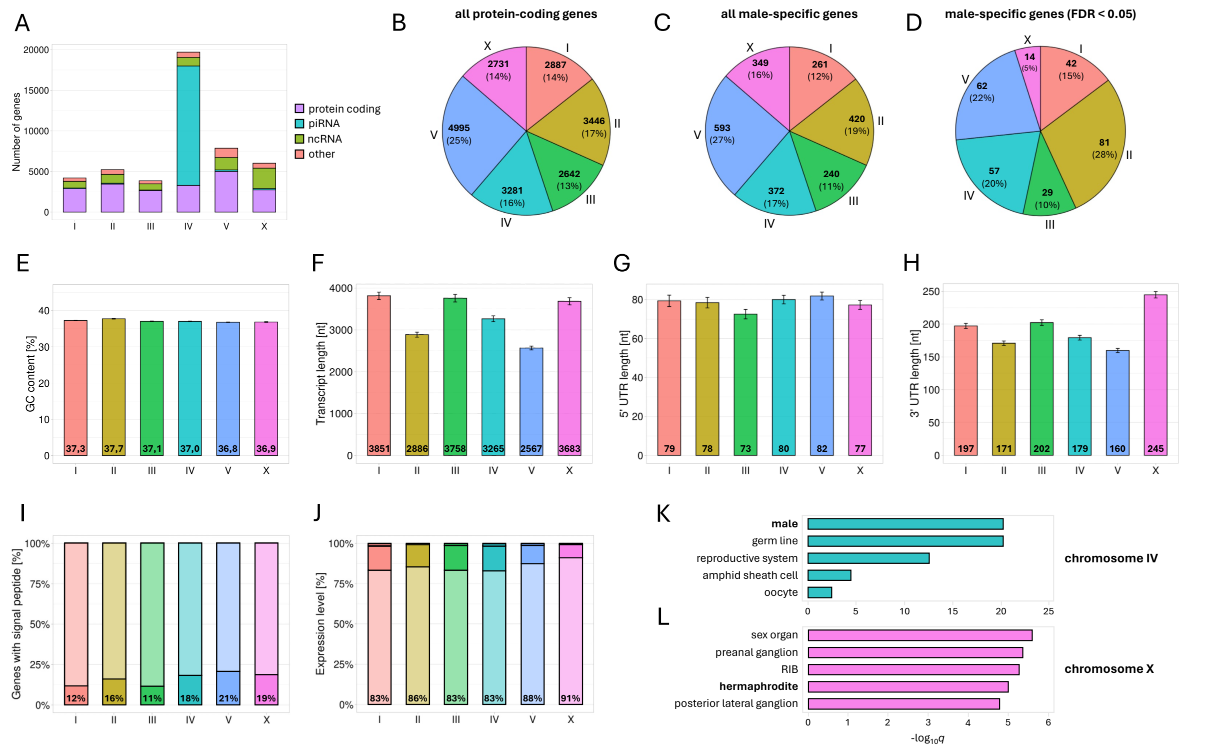


**Supplementary Figure S3. Differences in chromosome compositions.**

**(A)** Composition of each chromosome regarding RNA type. Four different types of RNA were highlighted: protein-coding mRNAs (purple), piRNA (cyan), ncRNA (green), and all other types (orange). The dataset for analysis was downloaded from ParaSite BioMart (WS291).

**(B-D)** Pie charts representing distributions of all protein-coding genes **(B)**, all male-specific genes **(C),** and significantly upregulated male-specific genes **(D)** between different chromosomes. The dataset for analyses was downloaded from ParaSite BioMart (WS291) and only protein-coding genes were included. Numbers on charts represent the number and the percentage of genes on each chromosome.

**(E)** Differences in GC content between genes localized on different chromosomes. The dataset for analysis was downloaded from ParaSite BioMart (WS291) and only protein-coding genes were included. Barplots represent mean values with SD.

**(F-H)** Differences in transcript length (unspliced transcript + UTRs) **(F)**, 5′ UTR length **(G),** and 3′ UTR length **(H)** between genes localized on different chromosomes. The dataset for analyses was downloaded from ParaSite BioMart (WS291) and only protein-coding genes were included. The average length of 5′ UTR and 3′ UTR is calculated considering the longer UTRs for each gene (the analyses were done for a narrowed-down subset of genes available in the dataset with necessary annotation). Barplots on each plot represent mean values with SD.

**(I)** Fractions of genes from each chromosome possessing signal peptide sequence based on studies by Suh J., Hutter H. (2012) (71). All genes detected in our DRS data were included. Numbers on the bars represent the percentage of genes with signal peptide sequence.

**(J)** Fractions of genes from each chromosome characterized by low, medium, or high expression levels. Expression levels are defined by baseMean values presented in Supplementary Table S1. Genes with baseMean < 20 are defined as lowly expressed, with 20 < baseMean < 500 as medium expressed, and with baseMean > 500 as highly expressed. All genes detected in our DRS data were included. Numbers on the bars represent the percentage of lowly expressed genes.

**(K)** and **(L)** Top five Tissue Enrichment terms for genes encoded on chromosome IV **(K)** or chromosome X **(L)** ordered by adjusted *p*-value (WormBase Enrichment Suite).

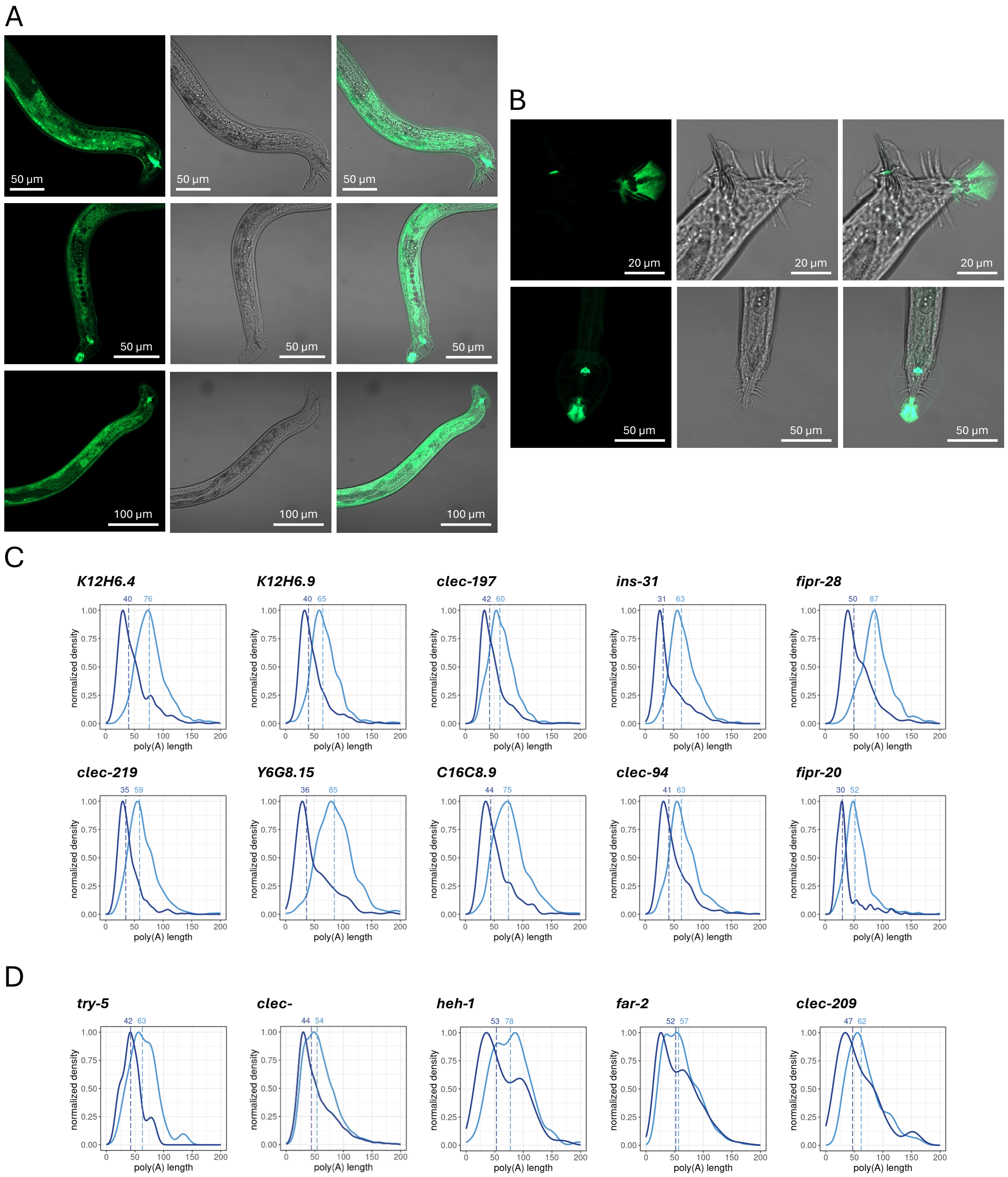


**Supplementary Figure S4. TENT-5 regulates the expression of male-specific genes.**

**(A)** Additional fluorescence microscopic images of TENT-5-GFP expression in *C. elegans* males (*tent-5(rtt6[tent-5::gfp::3xflag]* I). All images show the tail-end of a male body. Images were taken with a 40x lens and merged with bright-field images.

**(B)** Autofluorescence of the tail and copulatory spicules in wild-type (upper panel) and *tent-5:gfp* (lower panel) *C. elegans* males. Images were taken with a 40x lens and merged with bright-field images.

**(C)** Top ten protein-coding transcripts with the most significant poly(A) shortening in *tent-5(tm3504)* mutant males according to adjusted *p*-value (Supplementary Table S4). Poly(A) distributions are shown as light and dark blue for wild-type males and *tent-5(tm3504)* mutants, respectively. Vertical dashed lines represent the median poly(A) tail length for each condition (in nucleotides). The plots are normalized to 1.

**(D)** Density plots showing differences in the poly(A) tail distribution for genes or groups of genes mentioned individually in the chapter *"TENT-5 regulates the expression of male-specific genes"*. Vertical dashed lines represent the median poly(A) tail length for each condition (in nucleotides). The plots are normalized to 1.

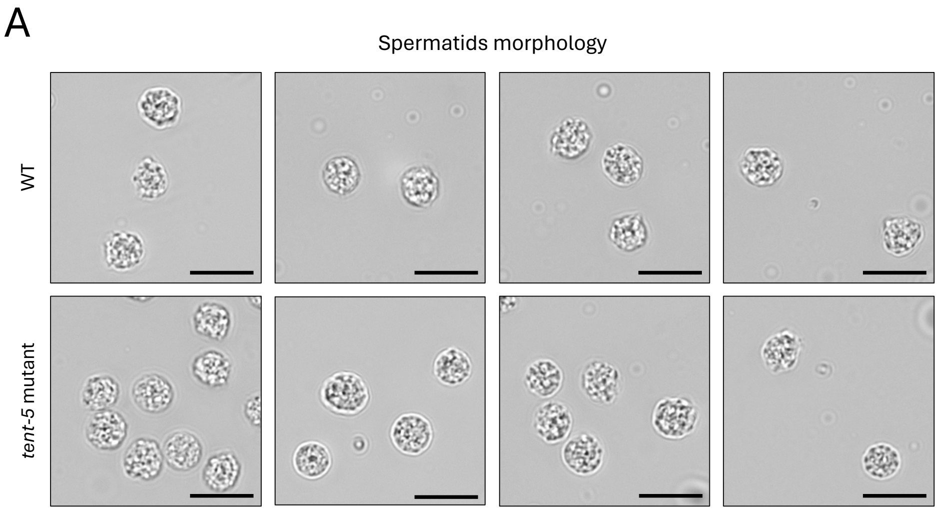


**Supplementary Figure S5. Comparison of spermatids morphology between wild-type and *tent-5* mutant males.**
**(A)** Additional microscopic images of spermatids morphology for wild-type and *tent-5* mutant males. Images are taken with 100x magnification. Scale bars => 10 microns.

**Supplementary Movie S1. Mating behavior experiment performed by crossing wild-type (N2) or *tent-5(tm3504)* mutant males with *unc-45* hermaphrodites.** The movie is shown at 20x speed.
